# Resurrected Rubisco suggests uniform carbon isotope signatures over geologic time

**DOI:** 10.1101/2021.05.31.446354

**Authors:** Mateusz Kedzior, Amanda K. Garcia, Meng Li, Arnaud Taton, Zachary R. Adam, Jodi N. Young, Betul Kacar

## Abstract

The earliest geochemical indicators of microbes—and the enzymes that powered them—extend back almost 3.8 billion years on our planet. Paleobiologists often attempt to understand these indicators by assuming that the behaviors of modern microbes and enzymes are consistent (uniform) with those of their predecessors. A uniformitarian assumption (i.e., the idea that fundamental geobiological processes have occurred in much the same manner over Earth history) seems at odds with our understanding of the inherent variability of living systems. Here, we examine whether a uniformitarian assumption for an enzyme thought to generate carbon isotope indicators of biological activity, RuBisCO, can be corroborated by independently studying the history of changes recorded within RuBisCO’s genetic sequences. Specifically, we resurrected a Precambrian-age, ancient RuBisCO by engineering its ancient DNA inside a modern cyanobacterium genome and measured the engineered organism’s fitness and carbon-isotope-discrimination profile. The envelope of ancestral RuBisCO isotopic fractionation observed here indicates that uniformitarian assumptions may be warranted, but with important caveats. Our results suggest that further inquiries that link molecule-level evolutionary changes with planet-level geochemical conditions are needed to discern whether enzyme-affected isotope fractionation trends extend deeper into the early Precambrian. Experimental studies illuminating life’s early molecular innovations are crucial to explore the foundations of Precambrian uniformitarian assumptions.

## Introduction

The history of life on Earth may be broadly subdivided into two, mutually exclusive macroevolutionary phases, the Phanerozoic and the Precambrian (Butterfield, 2007). The Phanerozoic (∼542 million years ago to present) may be characterized by physiological and anatomical innovations and their resultant effects on ecosystem expansion, trophic tiering, and sociality (Bambach, 1986; Bottjer and Ausich, 2016; Hannisdal and Peters, 2011). Hard- and soft-anatomical preservation provides a rich template for reconstructing Phanerozoic adaptive trends, correlating them with geographical and climatological changes (Sepkoski, 2015; Vermeij, 1973), and for testing observed diversity trends against possible systematic effects of preservation bias (Hendy, 2010; Peters, 2005; Peters and Foote, 2016). By contrast, the Precambrian (the ∼4 billion years preceding the Phanerozoic) is primarily characterized by genetic and metabolic biomolecular innovations, traded amongst microscopic organisms of uncertain phylogenetic assignment (Knoll, 2015; Schopf, 1994). The Precambrian record of evolutionary change appears to be cryptic and may have been comparatively static. This may be attributable to macroevolutionary dynamics that were distinctly non-Phanerozoic, or it may merely indicate a lack of direct paleontological and geological evidence of the specific timing and extent of intermediate biomolecular adaptive steps (Butterfield, 2007).

Comparative analyses of extant organisms have traditionally been the most informative means of interpreting the scant direct evidence of Precambrian life, but such analyses inevitably face pitfalls. A reasonable null hypothesis is that evolution is largely a uniformitarian process, such that rates or tempos may change but the primary modes of evolutionary change (as established through observations of the more comprehensive Phanerozoic record) are stable across geologic timescales (Erwin, 2011). Similarly, observations regarding present-day or recent geobiological and molecular processes might thus be relevant for interpretation of paleobiological signatures from the Precambrian. Such uniformitarian assumptions of ancient biology inferred from extant or Phanerozoic phenotypes are often employed to make sense of the Precambrian record, including body plan function (Adam et al., 2017; Yin et al., 2016), cladistic assignment (Butterfield, 2000; Golubic and Seong-Joo, 1999; Schopf, 2011; Tang et al., 2020), and isotope biosignature traces (Anbar and Rouxel, 2007; Beard, 1999; Garcia et al., 2021a; Schidlowski, 2001; Schopf et al., 2015). However, more recent studies question these assumptions, namely indicating that modes of biological variation can actually vary through time (Erwin, 2011). A major crux of the problem is that even the simplest modern organisms, as well as the macromolecules that compose them, differ from their ancient predecessors in having been shaped by the cumulative effects of billions of years of Earth-life co-evolution and ecosystem upheaval. For this reason, Precambrian functional or sequence comparisons may be of limited utility at organismal or biomolecular levels of adaptation, undercutting interpretations made possible by uniformitarian view. Novel experimental approaches may help to distinguish inferred paleobiological phenotypes from characteristically modern adaptive overprints.

The interpretation of the Precambrian carbon isotope record, comprising the oldest signatures of life on Earth, may be aided by novel experimental constraints on ancient phenotypes. Interpretations of this record is conventionally subject to uniformitarian assumptions regarding ancient biogeochemistry (Krissansen-Totton et al., 2015; Schidlowski, 2001). The ^13^C/^12^C isotopic mean of preserved organic carbon (δ^13^C_org_ ≈ -25‰) has remained notably static over geologic time (Garcia et al., 2021a; Krissansen-Totton et al., 2015; Schidlowski, 2001), and is leveraged as a general signature of ancient biological activity (Bell et al., 2015; Schidlowski, 1988; Schopf et al., 2018). Given its role in the Calvin-Benson-Bassham cycle— likely the predominant mode of carbon fixation for much of Earth history (Schidlowski, 2001)—the majority of contextual information used to assess how carbon isotope biosignatures might have been generated over Earth’s deep history comes from studies of the modern enzyme RuBisCO (Ribulose 1,5-Bisphosphate (RuBP) Carboxylase/Oxygenase, EC 4.1.1.39). RuBisCO catalyzes the uptake of inorganic CO_2_ from the environment and facilitates CO_2_ reduction and incorporation into organic biomass. The ^13^C/^12^C isotopic fractionation of modern Form I RuBisCO variants in photosynthetic organisms consistently measures ∼-25‰ (Guy et al., 1993; Roeske and O’Leary, 1984; Scott et al., 2007; von Caemmerer et al., 2014), approximately the same isotopic difference observed between inorganic and organic carbon in the Precambrian rock record. The carbon isotope discrimination of RuBisCO has therefore been presumed to have remained constant over the history of life.

Recent data, however, demonstrate that RuBisCO can produce significantly different carbon isotope signatures within organic matter in response to external factors, such as levels of atmospheric CO_2_ and/or O_2_ (Eichner et al., 2015; Freeman and Hayes, 1992; Hurley et al., 2021; Wilkes et al., 2018) or cellular carbon concentrating mechanisms, which affect the catalytic efficiency of RuBisCO (Flamholz et al., 2019; Studer et al., 2014; Tcherkez et al., 2006). Internal factors may also affect fractionation, such as single-point mutations that can alter the interaction between RuBisCO and CO_2_ (McNevin et al., 2007). The sensitivity of RuBisCO activity and fractionation to such factors is particularly significant given evidence for broad-scale trends in environmental conditions and biological innovations over the enzyme’s evolutionary history. These include the progressive oxygenation of the Earth atmosphere (Lyons et al., 2014), the evolution of novel carbon concentrating mechanisms (Badger, 2003), and the diversification of RuBisCO-hosting taxa. Thus, it seems unlikely that ancestral forms under ancient environmental conditions generated the same isotope fractionation signal as descendent homologs in modern organisms. Though some variability might be expressed by individual carbon isotopic measurements in the geologic record within a broader range, one might expect more frequent and significant excursions in the ∼-25‰ isotopic mean given the enormity of geobiological developments over the past >3 billion years (Garcia et al., 2021a). An experimental assessment that combines the phylogenetic history of RuBisCO with the study of intra- and extracellular conditions may provide a more insightful basis for comparing extant and Precambrian carbon isotope fractionation patterns.

The Form I clade of the RuBisCO phylogeny (and its macroevolutionary tractability afforded through green plant, algal and cyanobacterial fossil lineages) makes it an exemplary paleomolecular system for assessing uniformitarian assumptions applied to Precambrian biosignatures. Here, we establish an experimental system for the reconstruction of ancestral biomolecules with which to interpret evidence of Precambrian biological activity. Specifically, we report the resurrection and genetic incorporation of a phylogenetically reconstructed, ancient Form IB RuBisCO variant in a modern strain of cyanobacteria, *Synechococcus elongatus* PCC 7942 (thereafter *S. elongatus*) (Berla et al., 2013; Chen et al., 2012). We compared expression and activity levels of RuBisCO variants and the resulting changes in the growth of *S. elongatus* and isotope fractionation under ambient air as well as a CO_2_ concentration that reflects Precambrian conditions.

## Results

### Construction of a *S. elongatus* strain harboring ancestral RuBisCO

To experimentally investigate the generation of carbon isotope biosignatures in deep time, we designed a paleomolecular system to engineer computationally inferred, ancestral RuBisCO enzymes in extant cyanobacteria. We previously reconstructed a comprehensive phylogenetic history of RuBisCO and inferred maximum-likelihood ancestral RuBisCO large-subunit (RbcL) protein sequences (Kacar et al., 2017c) (Fig. 1A). For this study, we selected the ancestor of the Form IB RuBisCO clade (cyanobacteria, green algae, and land plants) for laboratory resurrection, designated “AncIB.” Chlorophyte and land plant RuBisCO homologs are nested among cyanobacterial sequences within the Form IB clade. The Form IB topology therefore recapitulates a primary plastid endosymbiotic history from cyanobacterial to Chlorophyte ancestors (Maruyama and Kim, 2020; Shih et al., 2016; Yoon et al., 2004) and constrains the minimum age of ancestral Form IB to older than the Archaeplastida. As a conservative estimate, AncIB is thus likely older than ∼1 Ga (the age of the oldest well-characterized, crown-group red and green algal fossils (Gibson et al., 2017; Tang et al., 2020)) and younger than maximum age estimates of cyanobacteria (∼3 Ga, as constrained by oxidized sediments potentially indicating the early presence of oxygenic photosynthesis (Anbar et al., 2007; Crowe et al., 2013)).

**Figure 1.**
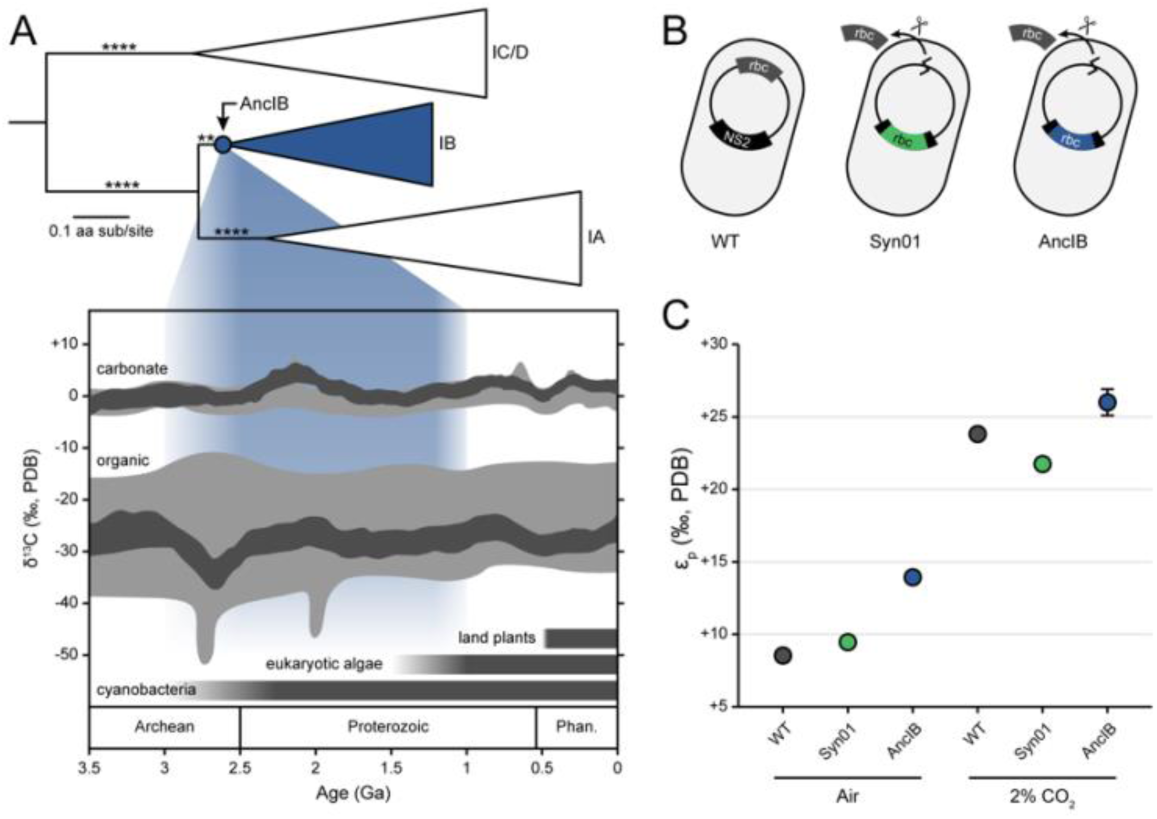
Reconstruction of ancestral RuBisCO biogeochemical signatures. (*A*) Maximum likelihood Form I RuBisCO RbcL phylogeny (derived from full RuBisCO phylogeny described in Kacar et al. (2017c)). Ancestral AncIB node and descendent Form IB clade highlighted in blue. Approximate likelihood ratio (aLR) branch support indicated by asterisks (**: >10, ****: >1000). Carbon isotope record figure adapted from Garcia et al. (Garcia et al., 2021a), with data from Schidlowski et al. (Schidlowski, 1988) (grey) and Krissansen-Totton et al. (Krissansen-Totton et al., 2015) (dark grey). Approximate age range of AncIB indicated by blue field (see text for discussion). (*B*) Genetic engineering of *S. elongatus* strains. Strain Syn01 was constructed by inserting a second copy of the *rbc* operon in the chromosomal neutral site 2 (NS2). Strain AncIB was constructed by inserting the genetic sequence encoding for the ancestral AncIB *rbcL* within the NS2 *rbc* operon. The native *rbc* operon was removed in both strains Syn01 and AncIB. (*C*) Photosynthetic carbon isotope fractionation (ε_p_) of *S. elongatus* strains in this study, cultured in ambient air or 2% CO_2_. Each data point represents the mean of three biological replicates and error bars indicate 1σ (error bars smaller than some datapoints).

The ancestral AncIB and *S. elongatus* native RbcL proteins differ at 37 sites and share 92% amino acid identity (Fig. 2). This site variation is evenly spread across the length of the protein, except for a highly conserved region between approximately site 170 to 285 (site numbering here and hereafter based on aligned WT *S. elongatus* RbcL; Fig. 2A) that constitutes a portion of the catalytic C-terminal domain and is proximal to the L-L interface and active site. Critical residues for carboxylase activity, including the Lys-198 site that binds CO_2_, are conserved in AncIB. Homology modeling of AncIB using the *S. elongatus* RbcL template (PDB: 1RBL (Newman et al., 1993)) indicates high structural conservation without predicted disruption to secondary structure (Fig. 2B). The nucleotide sequence for the reconstructed AncIB RbcL protein was codon-optimized for *S. elongatus* and cloned within a copy of the *rbc* operon into pSyn02 (Garcia et al., 2021b) for insertion into the *S. elongatus* chromosomal neutral site 2 (NS2). The native *rbc* operon was subsequently deleted to create the AncIB strain. In addition, we generated the control strain Syn01 harboring WT *rbcL* at NS2 (see Materials and Methods for full genome strategy). Thus, the AncIB and Syn01 strains carry a single ectopic copy of the *rbc* operon at NS2 and only differ in the coding sequence of the large RuBisCO subunit (Table S1).

**Figure 2.**
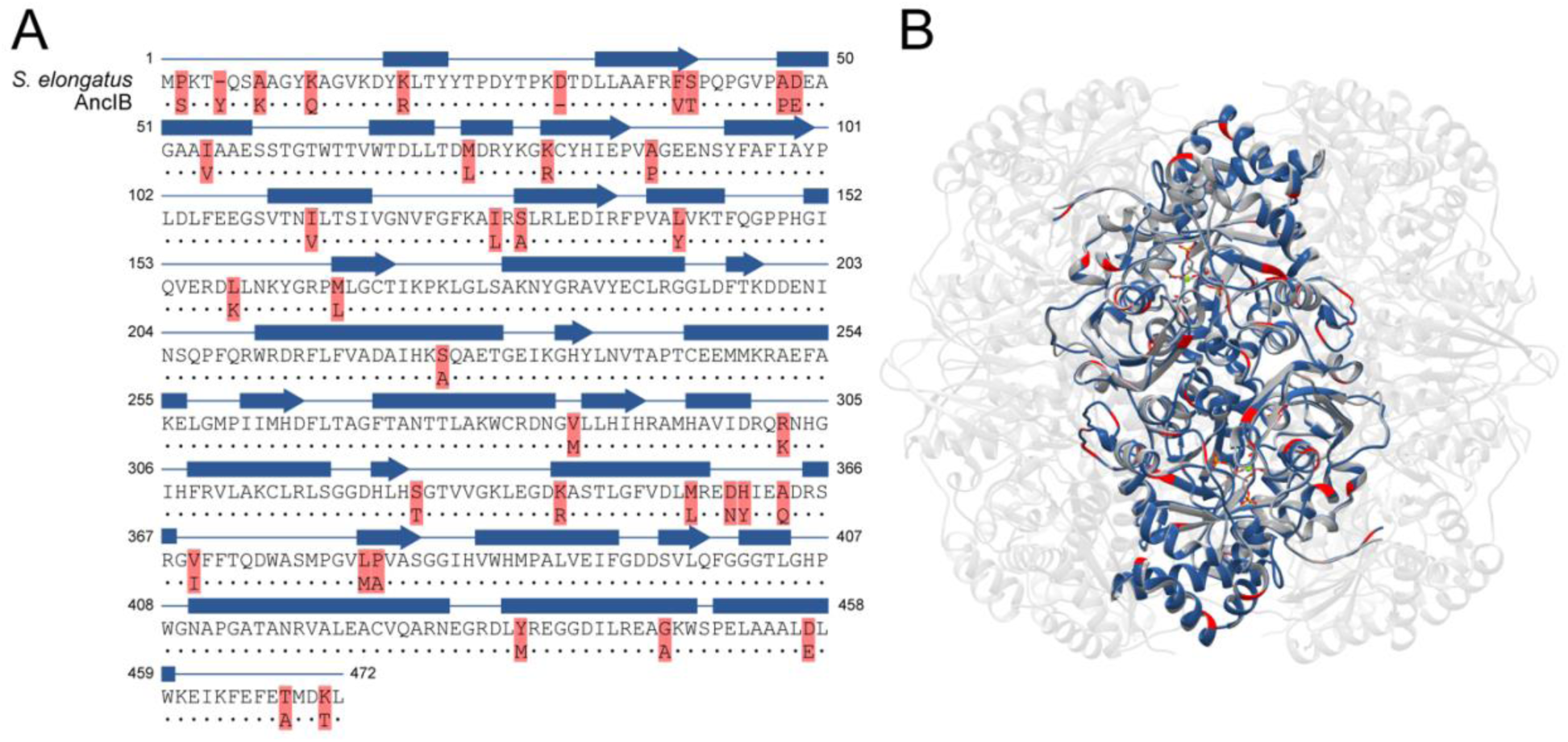
Structure and sequence features of ancestral RuBisCO. (*A*) Amino acid sequence alignment between ancestral AncIB and extant *S. elongatus* RbcL. Ancestral site variation relative to the *S. elongatus* template is highlighted in red. (*B*) Modeled structure of the ancestral AncIB L_2_ dimer (blue), aligned to the active conformation of the extant *S. elongatus* L_8_S_8_ hexadecamer (grey; PDB: 1RBL (Newman et al., 1993)). Highlighted residues in (*A*) are also highlighted in (*B*). Site numbering from extant *S. elongatus*. Conserved residues are indicated by dots and secondary structure is indicated above the sequences (blue rectangle: α-helix; blue arrow: β-sheet).

### Ancestral RuBisCO complements photoautotrophic growth of extant *S. elongates*

We cultured wild-type (WT) and engineered (AncIB and Syn01) strains of *S. elongatus* in both ambient air and 2% CO_2_ to evaluate the physiological impact of ancestral RuBisCO under estimated Precambrian CO_2_ concentrations (Catling and Zahnle, 2020) (Fig. 3A). The AncIB strain was capable of photoautotrophic growth in both ambient air and 2% CO_2_. Maximum growth rates for all strains were relatively comparable under each atmospheric condition (averaging doubling times of ∼15 to 20 hours), and generally increased under 2% CO_2_ relative to air (*p* < 0.001; Table S2). The AncIB strain exhibited a significantly diminished maximum cell density (OD_750_ ≈ 5) relative to both WT and Syn02 strains (OD_750_ ≈ 8; *p* < 0.001). No significant difference between WT and Syn01 growth rate or maximum cell density was observed under ambient air or 2% CO_2_. In addition, a moderate increase in the midpoint time for the AncIB strain was observed under air, indicating lag in growth (p < 0.01). These growth parameters taken together suggest decreased fitness of the AncIB strain relative to WT and Syn01.

**Figure 3.**
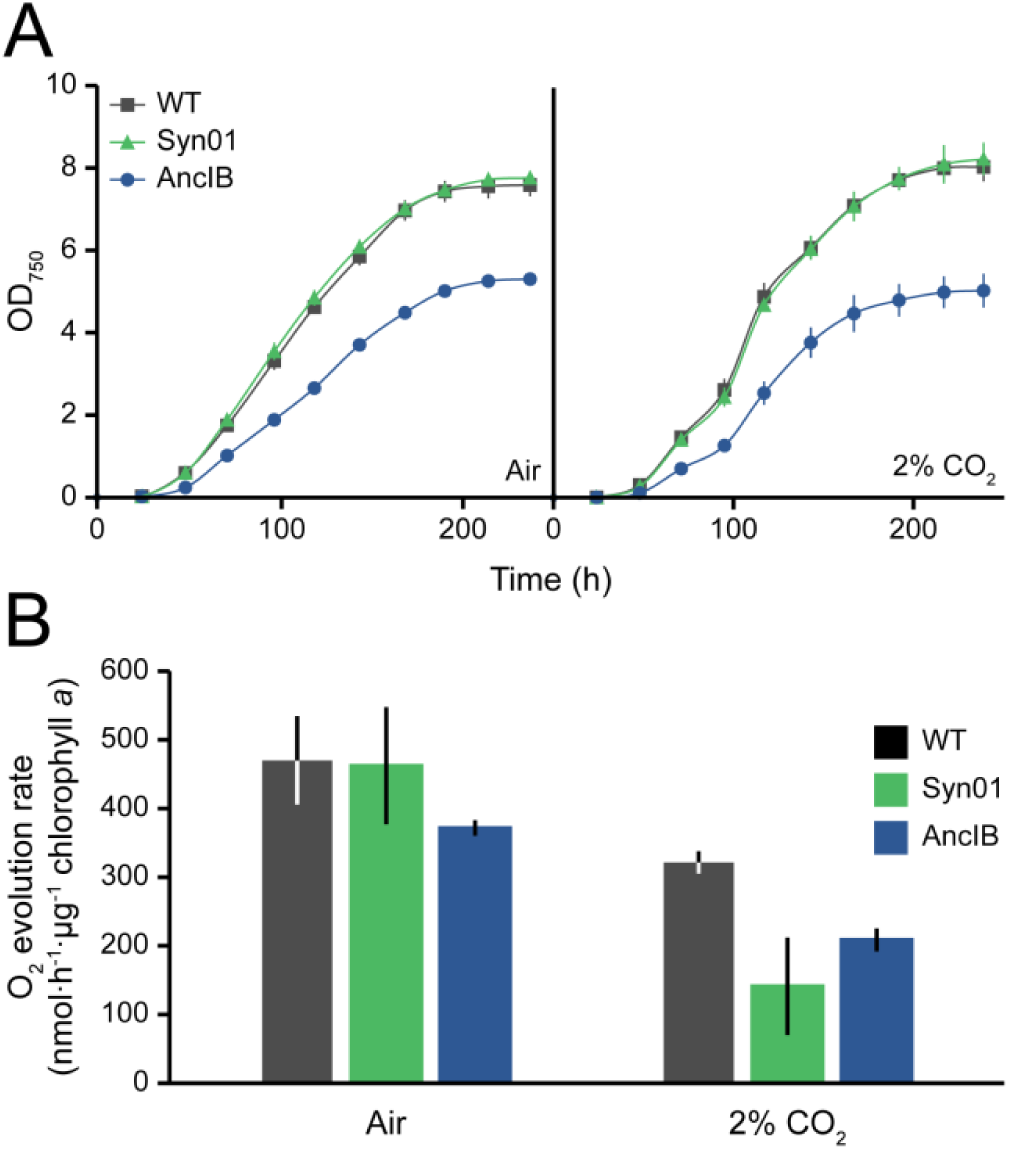
(*A*) Growth curves and (*B*) photosynthetic oxygen evolution of *S. elongatus* strains cultured in ambient air or 2% CO_2_. (*A, B*) Each data point or bar represents the mean of three biological replicates and error bars indicate 1σ.

AncIB RuBisCO protein produced a more modest impact on the oxygen evolution of *S. elongatus* compared to that on growth parameters. Cell suspensions were briefly incubated in the dark and subsequently exposed to saturated light in an electrode chamber to detect evolution of molecular oxygen (normalized to chlorophyll *a* concentration (Zavřel et al., 2015b)). For cells sampled from cultures grown in air, there was no statistical difference detected between any of the WT, Syn01, or AncIB strains (Fig. 3B). We did find a modest but significant decrease in photosynthetic activity for AncIB relative to WT for cells cultured in 2% CO_2_, generating ∼200 and ∼320 nmol O_2_·h^-1^·μg^-1^ chlorophyll *a*, respectively. However, a slower O_2_ evolution rate was also observed for Syn01 at 2% CO_2_.

### Ancestral RuBisCO is overexpressed and less catalytically active relative to WT

We assessed the impact of ancestral RuBisCO on gene expression at both the transcript and protein levels. *S. elongatus rbcL* transcript was measured by quantitative reverse-transcription PCR (RT-qPCR) and normalized to that of the *secA* reference gene (Szekeres et al., 2014). For strains cultured in ambient air, we found that the AncIB strain produced a ∼29-fold increase in *rbcL* transcript relative to WT or the control strain Syn01 (*p* < 0.001; Fig. 4A). The magnitude of AncIB *rbcL* overexpression was lower in 2% CO_2_, with a ∼4 and ∼2-fold increase observed relative to WT and the control strain, respectively (*p* < 0.001).

**Figure 4.**
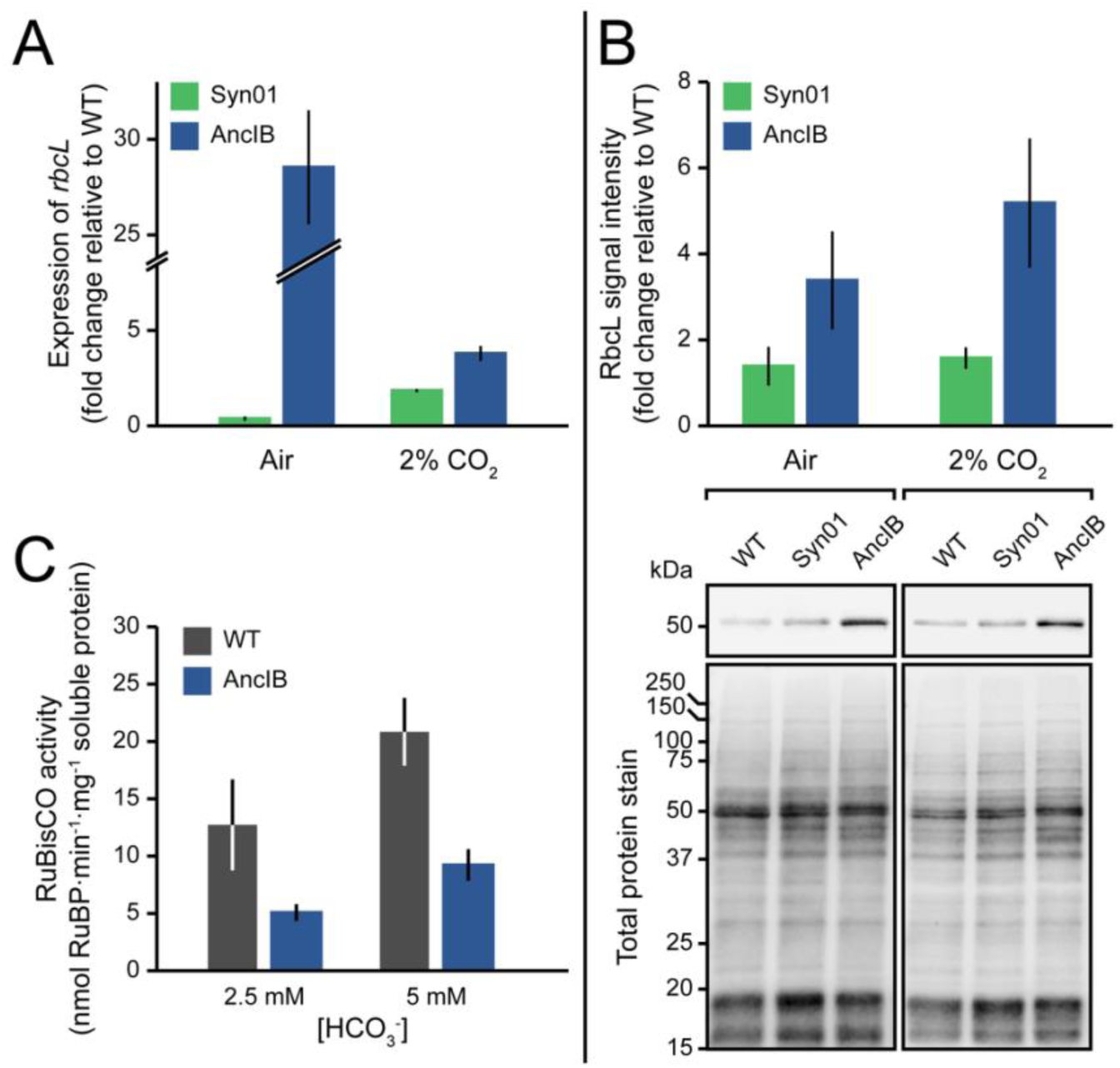
Expression and total cell activity of RuBisCO in *S. elongatus* strains. (*A*) Expression of *rbcL* in Syn01 and AncIB detected by RT-qPCR (*secA* reference gene), relative to WT. (*B*) Immunodetection of RbcL protein. *Top*, RbcL signal intensities normalized to those for total soluble protein load. *Bottom*, Western blot showing RbcL protein detected by anti-RuBisCO antibody and total protein stain from crude cell lysates (*C*) Total cell RuBisCO activity, measured with 2.5 mM and 5 mM HCO_3_^-^ concentrations. (*A-C*) Each bar represents the mean of three biological replicates (except the bar for 2% CO_2_ RbcL signal intensity data in (*B*), which represents the mean of six biological replicates) and error bars indicate 1σ.

RbcL protein was quantified for all *S. elongatus* strains by immunodetection using rabbit anti-RbcL antibody. We found that the amount of RbcL protein was also increased in the AncIB strain by ∼3-fold (*p* < 0.05) and ∼5-fold (*p* < 0.001) relative to WT or Syn01 in air and 2% CO_2_, respectively (Fig. 4B). No difference in RbcL quantity was detected between WT and Syn01 strains at either atmospheric condition. Finally, we confirmed assembly of the hexadecameric L_8_S_8_ RuBisCO complex in the AncIB strain by native PAGE and detection by anti-RbcL antibody (Fig. S1).

The total carboxylase activity of *S. elongatus* harboring ancestral RuBisCO was measured from crude cell lysates. Activity was assessed by an *in vitro* spectrophotometric coupled-enzyme assay that measures NADH oxidation and is reported as the RuBP consumption rate normalized to total soluble protein content (Kubien et al., 2008). For two sets of assays using either 2.5 mM or 5 mM HCO_3_ ^−^, AncIB lysate generated less than half the carboxylase activity of WT lysate (*p* < 0.05) (Fig. 4C).

### *S. elongatus* harboring ancestral RuBisCO produces greater carbon isotopic fractionation than wild-type

We measured the carbon isotope discrimination of *S. elongatus* strains cultured in ambient air and 2% CO_2_ to evaluate how the fractionation behavior of ancestral RuBisCO might influence the interpretation of ancient isotopic biosignatures preserved in the geologic record. The ^13^C/^12^C carbon isotope composition of *S. elongatus* was measured for biomass (δ^13^C_biomass_) as well as dissolved inorganic carbon (DIC; δ^13^C_DIC_) in the growth medium (Table S4). We calculated the carbon isotope fractionation associated with photosynthetic CO_2_ fixation (ε_p_) following Freeman and Hayes (Freeman and Hayes, 1992) after estimating δ^13^C_CO2_ from measured δ^13^C_DIC_ (Mook et al., 1974; Rau et al., 1996) (see Materials and Methods).

Overall, ε_p_ values were greater for *S. elongatus* strains cultured in 2% CO_2_ compared to ambient air. At 2% CO_2_, ε_p_ ranged between 21‰ and 26‰ compared to only 8‰ and 14‰ in air. We found that *S. elongatus* engineered with AncIB RbcL had an ε_p_ ∼5‰ greater than both WT and Syn01 when cultured in air (*p* < 0.001) and ∼2‰ to 4‰ greater than WT and Syn01 when cultured at 2% CO_2_ (*p* < 0.01) (Fig. 1C), though these differences in ε_p_ appear driven by the inorganic carbon pool composition rather than biomass (Table S4). A substantially smaller increase in ε_p_ (∼1‰; *p* < 0.001) was observed for Syn01 relative to WT under ambient air. Conversely, under 2% CO_2_, Syn01 ε_p_ was decreased by ∼2‰ (*p* < 0.001) relative to WT.

## Discussion

Form IB ancestral RuBisCO, when engineered into an extant strain of *S. elongatus*, decreased both the organismal growth capacity and the total cell RuBisCO activity. The genetic engineering strategy for insertion of the AncIB *rbcL* sequence in the cyanobacterial genome cannot solely account for these physiological differences since differences between the WT and Syn01 control strains were insignificant for most measured properties, or not comparable in magnitude to differences between the WT and AncIB strains. Rather, the observed differences for the AncIB strain indicate that the resultant phenotype is likely attributable to the functionality of the ancestral enzyme itself. The unique phenotype of the AncIB strain could be a direct result of the ancestral RbcL subunit or due to impediments to the assembly and activation of a hexadecamer RuBisCO complex containing both the ancestral RbcL and modern RbcS subunits, the latter of which was not targeted for ancestral reconstruction in this study. Our purposes here were to clarify the downstream physiological effects of a single ancestral RuBisCO gene, without potentially complicating factors arising from the interaction of two ancestral proteins. However, future studies could additionally reconstruct the Form IB RbcS and compare with the results presented here. Another possibility is hampered integration of ancestral RbcL given a modern suite of associated proteins required for RuBisCO folding and assembly (Gunn et al., 2020). Further, while overexpression of the ancestral RbcL occurred at the level of transcription and translation, the ancestral strain appears to have comparable levels of assembled hexadecamer RuBisCO, suggesting lower rates of RuBisCO assembly (or faster degradation). Even lower rates of measured total carboxylase activity suggest that the AncIB has decreased efficacy, which could be directly representing ancestral RuBisCO kinetics as well as the challenges associated with hybrid enzyme activation and activity.

Overexpression of the amount of ancestral RuBisCO shown by RT-qPCR and immunodetection assays is a common physiological response to decreased enzymatic efficacy throughout the cell (e.g., (Goldsmith and Tawfik, 2009; Kacar et al., 2017a)). However, expression compensation is insufficient to fully restore the extant WT phenotype, as indicated by the reduced fitness (i.e., decreased maximum cell density, oxygen evolution, and total carboxylase activity) of the ancestral strain harboring the ancestral RuBisCO compared to WT and Syn01.

There are few *in vitro* measurements of the kinetic isotope effect of Form IB RuBisCO in modern-day organisms, but those available range from ∼22‰ to 28‰ for cyanobacteria and C_3_ plants, respectively (Guy et al., 1993; Roeske and O’Leary, 1984; Scott et al., 2007; von Caemmerer et al., 2014). The ∼26‰ ε_p_ of AncIB strain biomass grown under 2% CO_2_ suggests that the ancestral RuBisCO also fractionates within this range. It has been theorized that RuBisCO kinetics have adapted in response to CO_2_ availability, either due to increased environmental CO_2_ (Tcherkez et al., 2006) or the emergence of CCMs (e.g., C_4_photosynthesis in plants (Christin et al., 2008)). Considering the positive relationship between enzymatic fractionation and RuBisCO’s specificity to CO_2_ (Tcherkez et al., 2006), reconstructing ancient RuBisCO kinetic isotope effect could provide insights into the co-evolution of atmospheric concentrations of CO_2_ and O_2_ and carbon fixation strategies during the Precambrian, in particular the emergence of carbon concentrating mechanisms (CCMs) (Burnap et al., 2015; Cameron et al., 2013; Hurley et al., 2021; Long et al., 2021). This is relevant as precise estimates of the magnitude of atmospheric CO_2_ elevation during the Precambrian relative to the present, as well as the emergence and effectiveness of Precambrian CCMs, are unknown.

We did observe statistically significant differences in ε_p_ of the ancestral strain compared to WT *S. elongatus* under both ambient air and 2% CO_2_ atmospheric conditions. However, upon inspection it appears that the diminished activity of the AncIB strain is influencing the composition of the DIC pool (both δ^13^C and concentration) in our cultures, and it is in fact the differences in DIC composition driving the calculated differences in ε_p_. Though strains were harvested at similar cell densities, small differences in cell concentrations at high densities can strongly influence carbonate chemistry of the media (Shi et al., 2009). The lower CO_2_ availability in the air treatment is more sensitive to cellular influence, resulting in a larger difference in ε_p_ compared to the 2% CO_2_ treatment. These differences in DIC are unlikely to be due to experimental setup (e.g., variations in CO_2_ bubbling), as biological replicates showed similar values. Therefore, the differences in fractionation reported here are likely implicated indirectly with the less efficient AncIB ancestral enzyme. Further comparative biomolecular characterization of AncIB and WT *S. elongatus* RbcL forms is needed to determine the degree to which enzymatic inefficiencies are contributing directly to the AncIB strain phenotype.

The observed carbon isotopic fractionation values corroborate a uniformitarian assumption for applying the maximal range of extant organism-level isotope fractionation values to interpret deep time isotopic biosignatures. The maximal observed isotopic fractionation of the AncIB strain under simulated Precambrian conditions with elevated CO_2_ is comparable to the ∼-25‰ isotopic mean of preserved, organic carbon across the plausible age range of the AncIB RuBisCO enzyme. These results suggest the conservation of RuBisCO fractionation behavior over the tested span of molecular evolution. There are, however, several potentially important contextual caveats for broader interpretation of these findings. The Form IB ancestor represents predecessors that are at least 1 billion years old, but it is also genetically and functionally still likely to be very different from the putative ‘root’ or common ancestor of all RuBisCO variants that emerged much earlier. With the observed *in vivo* functionality of AncIB RuBisCO reported here, we expect the successful reconstruction of older RuBisCO variants by similar means to be feasible. Reconstruction of older ancestors may further expand this maximal envelope of RuBisCO-generated carbon fractionation, or it may indicate that the extant maximal envelope is pervasive (and perhaps characteristic) across all functional variants of RuBisCO.

Another important caveat lies in the observation that, whereas all strains produce increased isotopic fractionation under elevated CO_2_, the comparative difference between ancestral AncIB and WT RbcL fractionation is relatively muted under 2% CO_2_ relative to ambient air. One possibility is that elevated CO_2_ brings the intrinsic fractionation properties of RuBisCO into relief (Bidigare et al., 1997; Hayes, 1993; Schubert and Jahren, 2012; Wilkes et al., 2018), at least compared to fractionation effects deriving from the overlying organismal physiology. By contrast, in present-day conditions, RuBisCO-mediated fractionation processes may be more significantly overprinted by physical factors that can affect RuBisCO catalytic efficiency, including cellular diffusion of O_2_/CO_2_ or other factors such as the presence of carbon concentration mechanisms.

There are many fundamental attributes of extant and ancestral metabolism for which the systemic effects on biosignature production have yet to be characterized. Disentangling these effects is critical for interpretation of the oldest biogeochemical record. A host cyanobacterium *S. elongatus* engineered with a Form IB RbcL ancestor confirms that organism- and enzyme-level effects on biosignature production are not always synonymous but differ in nuanced ways. These differences are contingent upon changes to internal (cellular, physiological) and external (environmental) conditions that have demonstrably varied over Earth’s long history. Cyanobacteria, with well-characterized genetic and morphological features (Berla et al., 2013; Burnap et al., 2015; Cameron et al., 2013; Chen et al., 2012; Long et al., 2021) and a tractable paleobiological history (Kacar et al., 2017b; Schopf, 2011), are ideal hosts for investigating a range of early Precambrian metabolic processes (Garcia and Kacar, 2019; Garcia et al., 2020; Kacar et al., 2017b).

Discernible trends (or steadfast consistencies) in metabolic outputs over macroevolutionary timescales can lead to foundational uniformitarian approaches to deep time molecular paleobiology. The available rock record becomes vanishingly sparse with greater age, but it is arguably well-sampled across key global-scale and biotically relevant isotopic systems at least through the early Archaean. Greater geologic sampling will therefore likely generate diminishing returns for shedding new light on deep time paleobiological trends. Innovative approaches that can chart a comprehensive envelope of biomolecular variability over time are a promising new means of reconciling coarse geochemical data with the nuance and complexity of ancient biological activity.

The engineering of ancient-modern hybrid organisms and their characterization can be used to complement the existing array of fossil remains, biogeochemical signatures, and modern organismal and molecular proxies to assess and contextualize plausible ranges of Precambrian carbon isotope biosignature production. Hybrid organisms may be particularly useful to disentangle the regulatory, physiological, and inter- and intramolecular factors that have impacted isotope fractionation, none of which are individually expressed in the geologic record. These factors must be systematically accounted for when interpreting bulk fractionation signals, even if only to elucidate the evolutionary molecular underpinnings of uniformitarian phenomena over geologic time.

After engineering a cyanobacterium with an ancient RuBisCO large protein subunit and cultivating it under conditions that mimic those prevailing through much of the Precambrian, we found the resultant carbon isotope fractionation to be within the range of organisms utilizing modern Form IB RuBisCO. The underlying biomolecular and organismal adjustments made by the cell to accommodate the ancestral gene were tracked, and we conclude that the small fractionation differences observed are likely attributable indirectly to decreased fitness of the AncIB strain, which influenced the inorganic carbonate chemistry of the media. The consistency of isotopic signatures generated by this strain indicates that uniformitarian assumptions based on the range of phenotypes of modern RuBisCO variants may apply for Precambrian environmental conditions, but that further study is warranted to discern organism- and enzyme-level trends in carbon isotope fractionation that may extend deeper into the early Precambrian.

### Limitations of the study

Our investigation characterized the *in vivo* attributes of a reconstructed, ancestral Form IB RuBisCO within a modern cyanobacterial to assess the commonly used, uniformitarian approach of interpreting ancient carbon isotope biosignatures based on comparable signatures generated by modern organisms and enzymes. Our results are supportive of the conservation of isotope fractionation behavior in cyanobacteria-hosted RuBisCO dating to the Precambrian. However, we do not provide an exhaustive assessment of all potential external factors that would together have shaped the isotope composition of preserved carbon. We target one time point in the evolution of RuBisCO using cyanobacteria as a model. Though the activity of RuBisCO enzymes have likely dominated primary production for billions of years, it has not been the only molecular determinant of the carbon isotope record (reviewed in (Garcia et al., 2021a)). A comprehensive exploration of the constancy of the carbon isotope trend across geologic timescales would require many detailed studies across environmental factors, including carbon fixation pathways catalyzed by different enzymes, and taxa that would have fluctuated in their proportional contributions to primary productivity over Earth history (e.g., prokaryotic versus eukaryotic contributions). Furthermore, our experimental system places an inferred, ancestral Form IB RuBisCO within the physiological context of a modern cyanobacterium. Such an approach is currently necessary to provide a tractable, experimental window into organism-level biogeochemical signatures stemming from the behavior of an ancestral enzyme. However, it is possible that the modern organism—including the associated modern proteins that interact with the ancestral RuBisCO protein—could influence the ancestral RuBisCO phenotype. These potential effects should in the future be investigated to provide more nuanced constraints on the range of isotopic biosignatures generated by ancient RuBisCO in the Precambrian. Our work here is a small start in the opening of an entirely new research direction that combines evolution, synthetic and molecular biology to experimentally evaluate the bases for interpreting the oldest record of life on Earth.

## Acknowledgements

We sincerely thank Emily Peñaherrera, Ryan Ward, and the University of California-Davis Stable Isotope Facility for assistance, and Jenan Kharbush for helpful discussions. We also thank two anonymous reviewers for their constructive comments and suggestions. This work was supported by the National Aeronautics and Space Administration Early Career Faculty (ECF) Award No. 80NSSC19K1617 (BK) and Interdisciplinary Consortium for Astrobiology Research No. 19-ICAR19_2-0007 (BK), the National Science Foundation Emerging Frontiers Program Award No. 1724090 (BK), National Aeronautics and Space Administration Postdoctoral Fellowship (AKG), Simons Foundation Early Career Award No. 561645 (JNY and ML), and National Institute of General Medical Sciences of the National Institutes of Health Award No. R01GM118815 (AT, to James W. Golden at the University of California-San Diego).

## Author contributions

Study design, and supervision, B.K.; methodology, M.K., A.K.G., M.L., A.T., J.Y., and B.K.; formal analysis, M.K. A.K.G. and B.K.; investigation, M.K., A.K.G., M.L., and B.K.; resources, A.T., J.Y., and B.K.; writing – original draft, M.K., A.K.G., Z.A., and B.K.; writing – review & editing, M.K., A.K.G., M.L., A.T., J.Y., Z.A., and B.K.; visualization, A.K.G. and B.K. All authors read, edited, and approved the final manuscript.

## Declaration of interests

The authors declare no competing interests.

## Star Methods

### Resource availability

### Lead contact

- Further information and requests for resources and reagents should be directed to and will be fulfilled by the lead contact, Betül Kaçar.

### Materials availability

- Strains and plasmids generated in this study are available from the lead contact, Betül Kaçar, upon request.

### Data and code availability

- All phylogenetic data has been deposited at GitHub (https://github.com/kacarlab/rubisco) and are publicly available as of the date of publication.
- This paper does not report original code.
- Any additional information required to reanalyze the data reported in this paper is available from the lead contact upon request.

## Experimental model and subject details

### Bacterial strains

*S. elongatus* PCC 7942 strains were cultured in BG-11 medium (Rippka et al., 1979) as liquid cultures or on agar plates (1.5% (w/v) agar and 1 mM Na_2_S_2_O_3_·5H_2_O). Liquid cultures were grown at 30°C, continuous shaking at 120 rpm, sparged with ambient air or 2% CO_2_, and with 115 µmol photon·m^-2^·s^-1^ (except for cultures used to prepare samples for O_2_ evolution, which were grown with 80 µmol photon·m^-2^·s^-1^). The 2% CO_2_ gas mix was controlled by an environment chamber (Percival, Cat. No. I36LLVLC8) with a CO_2_ tank input. For recombinant strains, liquid and solid media were supplemented with appropriate antibiotics: 2 μg·ml^−1^ Spectinomycin (Sp) plus 2 μg·ml^−1^ Streptomycin (Sm), 5 μg·ml^−1^ Kanamycin (Km). Cyanobacterial growth was measured at an optical density of 750 nm (OD_750_) and growth parameters were estimated using the Growthcurver package for R (Sprouffske and Wagner, 2016). Cultures were sampled at the middle exponential growth phase, i.e., at an OD_750_ of ∼2.5 (AncIB) or ∼4.5 (WT and Syn01) for all subsequent experiments.

## Method details

### Inference of ancestral AncIB RbcL protein

A RuBisCO RbcL phylogeny was reconstructed as previously described (Kacar et al., 2017c). Briefly, RbcL orthologs were identified from the NCBI protein database by BLAST (sequence dataset and the tree can be found at https://github.com/kacarlab/rubisco). Phylogenetic analysis was performed by Phylobot (Hanson-Smith and Johnson, 2016), a web portal that integrates alignment, phylogenetic reconstruction by RAxML (Stamatakis, 2014), and ancestral sequence inference by PAML (Yang, 2007). A maximum likelihood phylogeny was built using a MSAProbs alignment (Liu et al., 2010) and the best-fit PROTCATWAG model (Lartillot and Philippe, 2004; Whelan and Goldman, 2001), determined by the Akaike information criterion (Abascal et al., 2005). Ancestral states were reconstructed at each amino acid site for all phylogenetic nodes, and gap characters were inferred according to Fitch’s parsimony (Fitch, 1971).

### Cyanobacteria genetic engineering

Recombinant strains of *S. elongatus* were constructed by natural transformation using standard protocols (Clerico et al., 2007) with minor modifications (Garcia et al., 2021b). The plasmids and strains used in this study are listed in Table S1. The construction of *S. elongatus* Syn01, carrying a single ectopic copy of the *rbc* operon at NS2, as well as the plasmids pSyn01 and pSyn02 used to construct strain Syn01, were described previously (Garcia et al., 2021b). The construction of strain AncIB was performed similarly to Syn01. Briefly, pSyn03, which carries the AncIB nucleotide sequence within the entire *rbc* operon (including flanking sequences and homologous regions for recombination at neutral site 2 (NS2) of the *S. elongatus* chromosome), was transformed in *S. elongatus*. Transformation of WT *S. elongatus* with pSyn03 generated strain Syn03 carrying a second copy of the *rbc* operon at NS2. Strain Syn03 was subsequently transformed with pSyn01 to replace the native *rbc* operon with a spectinomycin/streptomycin resistance gene as described previously (Garcia et al., 2021b), producing strain AncIB. Transformants of Syn03 and AncIB were screened for complete segregation by colony PCR using primers F06, R06, F07, and R07 (Fig. S2; Table S3) and the strain sequences at the deletion and insertion sites were further verified by Sanger sequencing using the primers R07, F08, R08, F15, and F16 (Table S3). To construct plasmid pSyn03, pSyn02 excluding the *rbcL* coding sequence was PCR-amplified and linearized using primers F13/R13 and assembled with the AncIB *RbcL* coding sequence codon-optimized for *S. elongatus* and synthesized by Twist Bioscience. Both DNA fragments were assembled using the GeneArt™ Seamless Cloning and Assembly Kit (Invitrogen, Cat. No. A13288). (Table S3).

### RT-qPCR analysis of *rbcL* expression

Cells were pelleted by centrifugation and resuspended in TE buffer (10 mM Tris, pH 8.0, 1 mM EDTA). Total RNA was extracted using the RNeasy® Protect Bacteria Mini Kit (QIAGEN, Cat. No. 74524). DNase I-treated RNA was then used in reverse transcription (RT) performed with the SuperScript™ IV First-Strand Synthesis System (Invitrogen, Cat. No. 18091050). F09/R09, F11/R11, F14/R14 pairs of qPCR primers (Table S3) were designed with Primer3Plus (Untergasser et al., 2007) (http://www.bioinformatics.nl/cgi-bin/primer3plus/primer3plus.cgi). The quality of cDNA and primer specificity was assessed by PCR using cDNA templates (RT positive reactions) and RT negative controls. qPCR was performed by the real-time thermal cycler qTOWER^3^ G (Analytik Jena AG) using qPCRsoft software. The relative expression of native and AncIB *rbcL* was calculated as the average fold change normalized to the *secA* reference gene (Szekeres et al., 2014) using the delta-delta Ct method. The experiment was carried out using three biological replicates and three technical replicates.

### RbcL protein immunodetection

Cells were pelleted by centrifugation and resuspended in 95°C TE buffer supplemented with 1% (w/v) SDS and incubated at 95°C for 10 min. The mixture was sonicated and centrifuged to remove cell debris. Total protein concentration in the crude cell lysates was measured using the Pierce™ BCA Protein Assay Kit (Thermo Scientific, Cat. No. 23225). Lysates containing 5 μg of total protein in Laemmli sample buffer were loaded onto a 6% (v/v) polyacrylamide stacking gel. Proteins were electrophoresed in a 12% polyacrylamide resolving gel and blotted onto a nitrocellulose membrane. Detection of RbcL and total protein load was performed as previously described (Garcia et al., 2021b; Kędzior and Kacar, 2021). Detection of RbcL was performed with rabbit anti-RbcL antibody (Agrisera, Cat No. AS03 037) and rabbit IgG secondary antibody (LI-COR Biosciences, Cat No. 926-32211). The densitometric analysis of RbcL signal intensity, normalized to total protein load, was performed with Quantity One® software (Bio-Rad) for three to six biological replicates.

### RuBisCO assembly confirmation

Assembly of the RuBisCO large and small subunits into a hexadecameric complex in each strain was evaluated by native gel electrophoresis and immunodetection, as previously described (Garcia et al., 2021b; Kędzior and Kacar, 2021). Immunodetection of the RuBisCO complex was performed for three biological replicates with the same primary and secondary antibodies that were used to detect RbcL, as described above.

### RuBisCO catalytic activity

The activity of RuBisCO in cyanobacterial lysates was measured using a spectrophotometric coupled-enzyme assay that links this activity with the rate of NADH oxidation (Kubien et al., 2011). Cell lysis and the activity assay were carried out as previously described (Garcia et al., 2021b) with either 2.5 mM or 5 mM NaHCO_3_. After 20 min at 25 °C for activation of Rubisco, the reaction was initialized with the addition of ribulose 1,5-bisphosphate (RuBP) (0.5 mM) and the absorbance at 340 nm was recorded using a Synergy H1 plate reader (BioTek). RuBisCO activity was reported as the RuBP consumption rate normalized to total soluble protein content. The assay was performed for three biological replicates.

### Photosynthetic oxygen evolution rate

*S. elongatus* strain photosynthetic activity was assayed using a Clark-type oxygen electrode chamber to measure the level of molecular oxygen produced in cyanobacterial cultures. Cells were pelleted and resuspended in fresh BG-11 to an OD_750_ of ∼1 following De Porcelinis (De Porcellinis et al., 2018). Concentration of chlorophyll *a* (for normalization) was measured following the protocol by Zavrel et al. (Zavřel et al., 2015a). The remaining suspension was incubated in the dark for 20 min with gentle agitation. Samples from each suspension were analyzed in an oxygen electrode chamber under saturated light, using the Oxygraph+ System (Hansatech Instruments) equipped with the OxyTrace+ software. Oxygen evolution rate was monitored for 10 min and expressed as nanomoles of molecular oxygen evolved per hour per microgram of chlorophyll *a*. The assay was performed for three biological replicates.

### Carbon isotope fractionation

Cells were pelleted by centrifugation and washed in 10 mL of 10 mM NaCl (OD_750_ for Syn-1 ∼ 4.5, OD_750_ for AncIB ∼2.5). Pellets were then dried at 50°C. In parallel, the supernatant from centrifuged culture samples was sterilized through 0.2 µm filtration for DIC isotopic analysis of growth media. Sterilized media was transferred to Exetainer vials leaving no headspace and stored at 4°C until analysis. Isotopic analysis was performed for three biological replicates.

The carbon isotope composition of bulk biomass (δ^13^C_biomass_) and DIC (δ^13^C_DIC_) was determined at the UC Davis Stable Isotope Facility. δ^13^C_biomass_ was analyzed using a PDZ Europa ANCA-GSL elemental analyzer interfaced to a PDZ Europa 20-20 isotope ratio mass spectrometer (Sercon Ltd.). DIC samples were analyzed by gas evolution and composition was measured by a GasBench II system interfaced to a Delta V Plus IRMS (Thermo Scientific). The carbon isotopic composition values were reported relative to the Vienna PeeDee Belemnite standard (V-PDB):

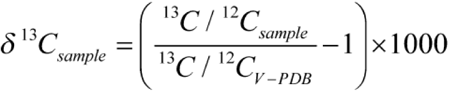

The isotopic composition of dissolved molecular CO_2_ (δ^13^C_CO2_) was estimated from δ^13^C_DIC_ following Rau et al. (Rau et al., 1996) and Mook et al. (Mook et al., 1974):

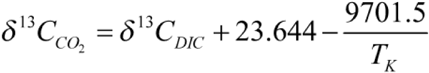

The carbon isotope fractionation associated with photosynthetic CO_2_ fixation (ε_p_) was calculated relative to δ^13^C_CO2_ in the post-culture medium according to Freeman and Hayes (1992):

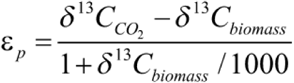

### Quantification and statistical analysis

Results for experimental analyses were presented as the mean and the sample standard deviation (1σ) values of at least three biological replicates. For comparisons of two groups, statistical significance was analyzed by an unpaired, two-tailed *t*-test assuming equal variance. For comparisons of three or more groups, significance was analyzed by one-way ANOVA and a post-hoc Tukey HSD test.

### Key resources table

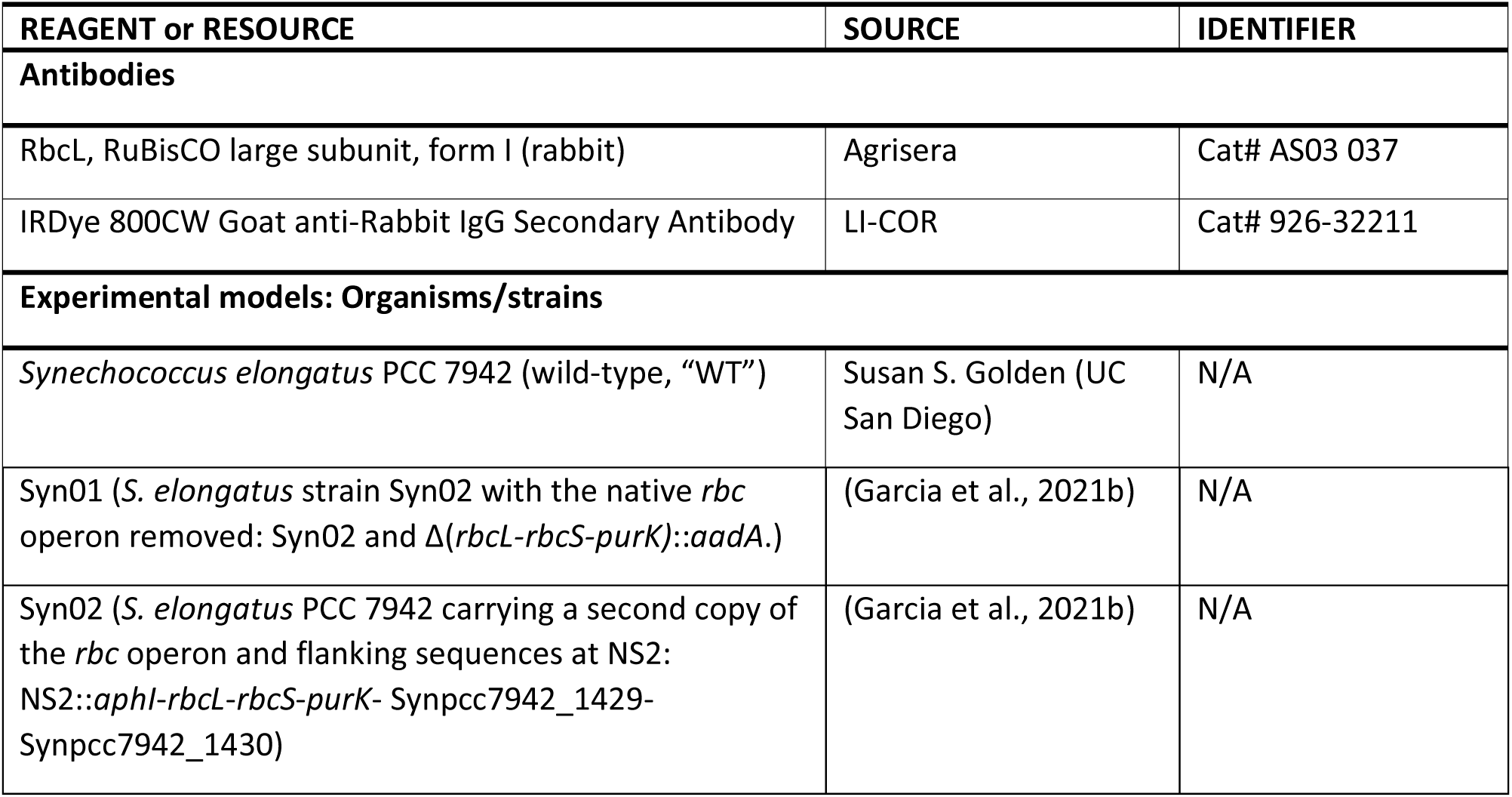
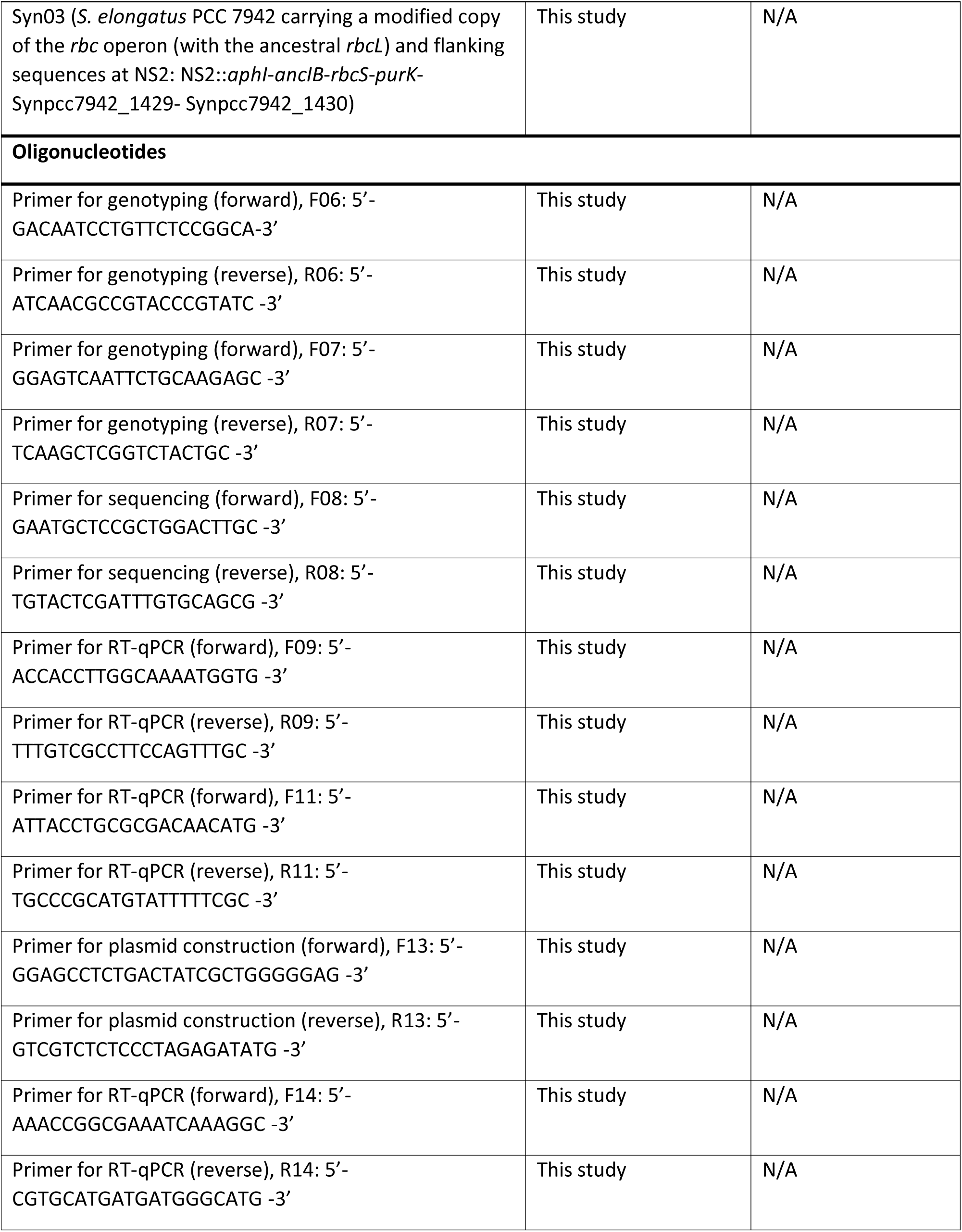
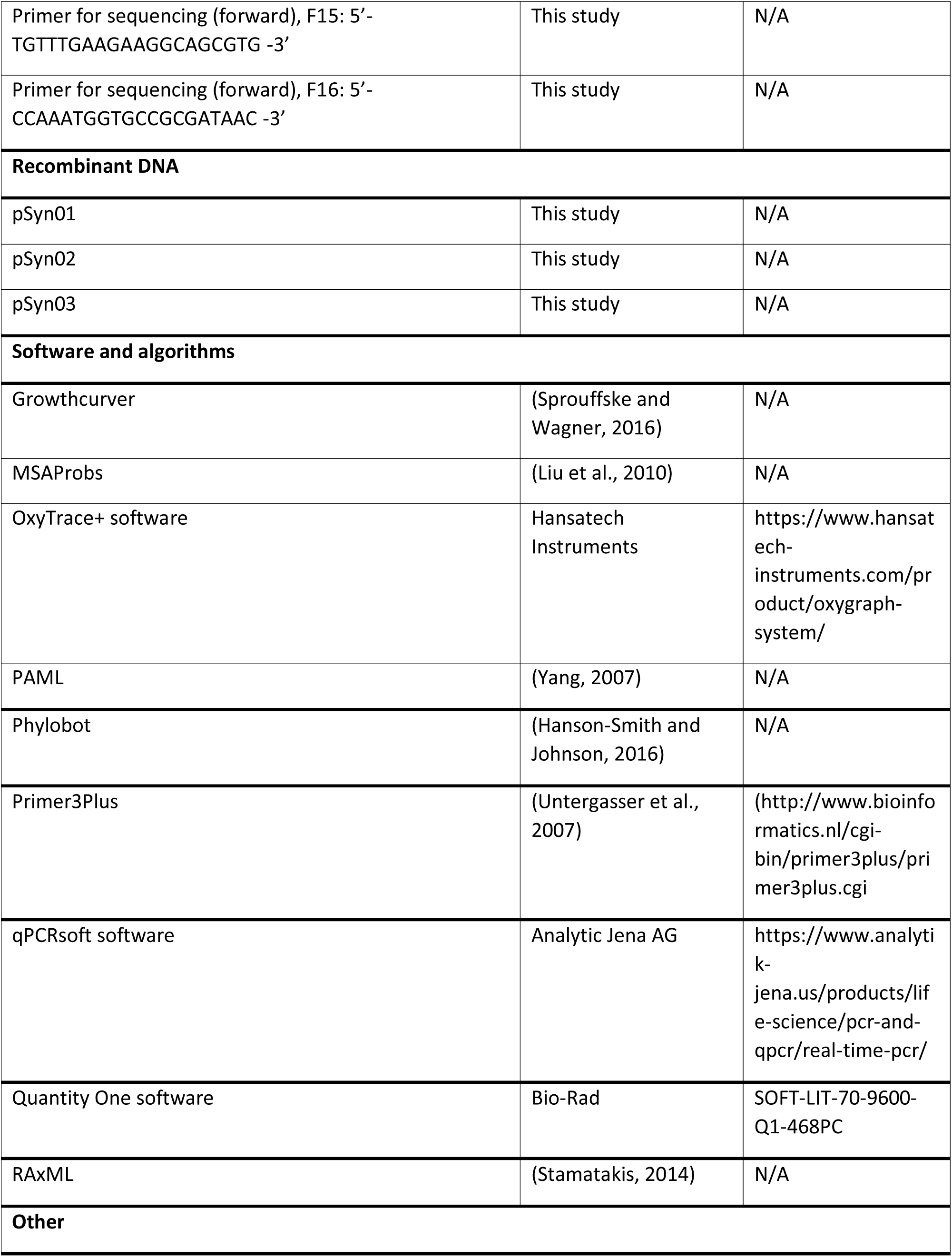
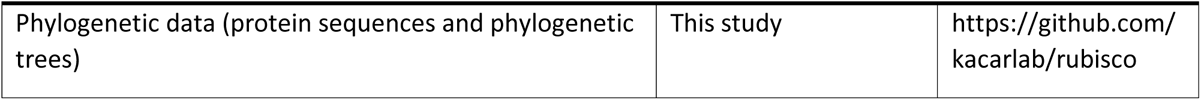

## Notes

### Competing Interest Statement

The authors have declared no competing interest.

### Summary of Updates

Updated the abstract, results and discussion. Added missing citations and updated figures.

