## Supplementary Information for "Resurrected Rubisco suggests uniform carbon isotope signatures over geologic time"

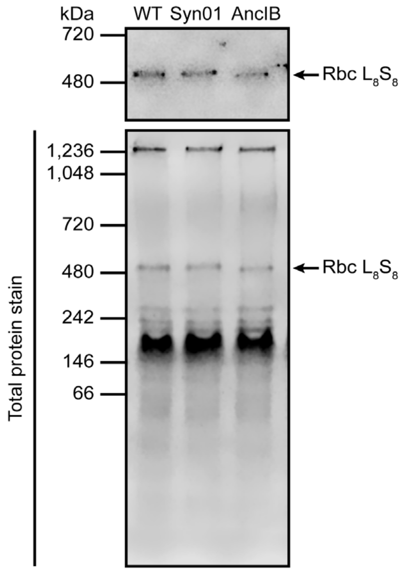


**Figure S1.** Immunodetection of assembled RuBisCO. Related to Figure 4. Western blot showing the assembly of RbcL (WT and AncIB) and RbcS into the $L_8S_8$ hexadecameric complex (520 kDa), detected by anti-RbcL antibody.

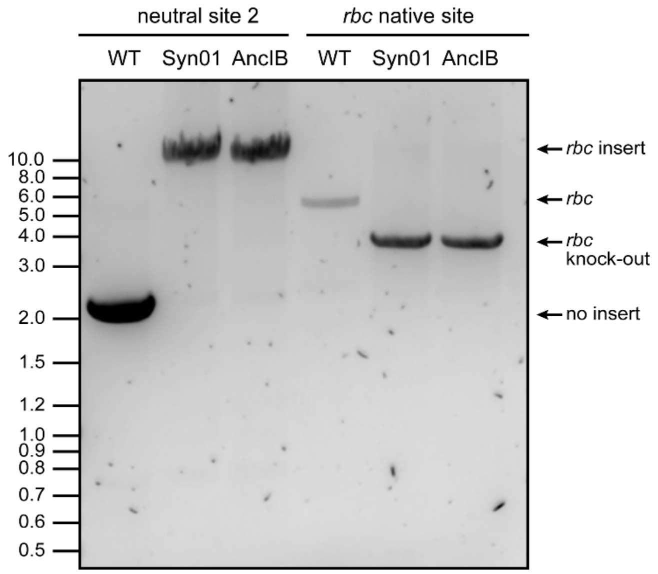


**Figure S2.** Genotyping of *S. elongatus* strains. Related to Figures 1 and 4. Primers F06/R06 (Table S3) were used to confirm the *rbc* operon insertion at the neutral site 2 (NS2), either with WT *rbcL* (for Syn01) or AncIB *rbcL* (*rbc* insert: 8,617 bp; no insert: 1,970 bp). Primers F07/R07 were used to confirm the presence or absence of the rbc operon at the native site (*rbc* present at its native site: 5,140 bp; *rbc* knocked out and replaced with the aadA gene: 3,342 bp).

**Table S1.** Strains and plasmids used in this study.

| Strain or plasmid | Description/Genotype | Antibiotic resistance | Source/Reference |
|---|---|---|---|
| WT | Wild-type strain of *S. elongatus* PCC 7942 | - | Susan S. Golden (UC San Diego) |
| Syn01 | *S. elongatus* strain Syn02 with the native *rbc* operon removed: Syn02 and Δ(*rbcL-rbcS-purK)*::*aadA*. | Km, Sp+Sm | (Garcia et al., 2021b) |
| Syn02 | *S. elongatus* PCC 7942 carrying a second copy of the *rbc* operon and flanking sequences at NS2: NS2::*aphI-rbcL-rbcS-purK*- Synpcc7942_1429-Synpcc7942_1430 | Km | (Garcia et al., 2021b) |
| Syn03 | *S. elongatus* PCC 7942 carrying a modified copy of the *rbc* operon (with the ancestral *rbcL*) and flanking sequences at NS2: NS2::*aphI-ancIB-rbcS-purK*- Synpcc7942_1429-Synpcc7942_1430 | Km, Sp+Sm | This study |
| AncIB | *S. elongatus* strain Syn03 carrying the ancestral AncIB *rbcL* gene in the *rbc* operon copy at NS2 and having the native *rbc* operon replaced with a Sp/Sm resistance gene: Δ(*rbcL-rbcS-purK)*::*aadA*. | Km | This study |
| pSyn01 | Plasmid to replace *S. elongatus'* native *rbc* operon (CP000100: 1479070-1482595) with a Sp/Sm resistance gene: Δ(*rbcL-rbcS-purK)*::*aadA*. | Sp+Sm | (Garcia et al., 2021b) |
| pSyn02 | Plasmid for recombination at NS2 of *S. elongatus* chromosome carrying the *rbc* operon including *rbcL*, *rbcS*, *purK,* and flanking sequences from *S. elongatus* PCC 7942 (CP000100: 1479071-1484283) | Km | (Garcia et al., 2021b) |
| pSyn03 | pSyn02 in which the coding sequence of RbcL was replaced with the coding sequence of AncIB | Km | This study |

**Table S2. Growth parameters of *S. elongatus* strains.** Values represent the mean of three replicates ± 1σ. Asterisks indicate significance relative to WT for the same atmospheric condition, determined by one-way ANOVA and post-hoc Tukey HSD tests (**: $p < 0.01$; ***: $p < 0.001$).

| Strain | Atmosphere | Doubling time (h) | Midpoint time (h) | Maximum cell density ($OD_{750}$) |
|---|---|---|---|---|
| WT | Air | 18.9 ± 1.0 | 106.4 ± 2.8 | 7.7 ± 0.3 |
| Syn01 | | 18.6 ± 0.3 | 103.8 ± 1.9 | 7.7 ± 0.1 |
| AncIB | | 20.5 ± 0.4 | 118.4 ± 2.1** | 5.4 ± 0.2*** |
| WT | 2% $CO_2$ | 16.9 ± 1.5 | 110.8 ± 6.1 | 8.0 ± 0.3 |
| Syn01 | | 17.3 ± 1.0 | 113.9 ± 2.3 | 8.1 ± 0.5 |
| AncIB | | 15.8 ± 0.5 | 117.8 ± 1.7 | 5.0 ± 0.4*** |

**Table S3.** Primers used in this study.

| **Primer**[a] | **Sequence (5'-3')** | **Description** |
|---|---|---|
| **F06** | GACAATCCTGTTCTCCGGCA | Genotyping *S. elongatus* strains to screen for the *rbc* operon insert (either with native or AncIB *rbcL*) at NS2 (PCR product size: with *rbc* insert – 8,617 bp, without *rbc* insert – 1,970 bp). |
| **R06** | ATCAACGCCGTACCCGTATC | |
| **F07** | GGAGTCAATTCTGCAAGAGC | Genotyping *S. elongatus* strains to confirm the presence or absence of the *rbc* operon at the native site (PCR product size: with the *rbc* operon – 5,140 bp, without the *rbc* operon – 3,342 bp) R07 was also used to sequence the native *rbc* operon deletion site. |
| **R07** | TCAAGCTCGGTCTACTGC | |
| **F08** | GAATGCTCCGCTGGACTTGC | Sequencing the *rbc* operon insertion site at NS2. F08 was also used to sequence the native *rbc* operon deletion site. |
| **R08** | TGTACTCGATTTGTGCAGCG | |
| **F09** | ACCACCTTGGCAAAATGGTG | qPCR analysis of the expression of *rbcL* (ID: Synpcc7942_1426) that encodes the RuBisCO large subunit. |
| **R09** | TTTGTCGCCTTCCAGTTTGC | |
| **F11** | ATTACCTGCGCGACAACATG | qPCR analysis of the expression of *secA* (reference gene, ID: Synpcc7942_0289) that encodes the preprotein translocase subunit SecA. |
| **R11** | TGCCCGCATGTATTTTTCGC | |
| **F13** | GGAGCCTCTGACTATCGCTGGGGGAG | Linearization of pSyn02 without the *rbcL* coding sequence. |
| **R13** | GTCGTCTCTCCCTAGAGATATG | |
| **F14** | AAACCGGCGAAATCAAAGGC | qPCR analysis of the expression of AncIB *rbcL* that encodes the AncIB RuBisCO large subunit. |
| **R14** | CGTGCATGATGATGGGCATG | |
| **F15** | TGTTTGAAGAAGGCAGCGTG | Sequencing the internal region of AncIB *rbcL* inserted at NS2. |
| **F16** | CCAAATGGTGCCGCGATAAC | |

[a]F - forward, R – reverse

**Table S4.** Isotopic composition of biomass and DIC in growth medium of *S. elongatus* cultures.

| Strain | Atmosphere | $\delta^{13}C_{biomass}$ (‰) | $\delta^{13}C_{DIC}$ (‰) | $\delta^{13}C_{CO2}$ (‰) | [DIC] (mM) |
|---|---|---|---|---|---|
| WT | Air | -19.63 ± 0.07 | -2.92 ± 0.02 | -11.28 ± 0.02 | 5.14 ± 0.07 |
| | 2% $CO_2$ | -26.41 ± 0.15 | 5.77 ± 0.30 | -2.59 ± 0.30 | 7.35 ± 0.24 |
| Syn01 | Air | -20.20 ± 0.19 | -2.63 ± 0.10 | -11.00 ± 0.10 | 5.23 ± 0.02 |
| | 2% $CO_2$ | -26.84 ± 0.10 | 3.29 ± 0.25 | -5.07 ± 0.25 | 7.53 ± 0.34 |
| AncIB | Air | -20.60 ± 0.08 | 1.26 ± 0.10 | -7.10 ± 0.10 | 3.44 ± 0.05 |
| | 2% $CO_2$ | -31.2 ± 0.34 | 3.15 ± 0.62 | -5.21 ± 0.62 | 5.70 ± 0.70 |
